# Aging amplifies sex differences in low alpha and low beta EEG oscillations

**DOI:** 10.1101/2024.07.31.603949

**Authors:** Chuanliang Han, Vincent C.K. Cheung, Rosa H.M. Chan

## Abstract

Biological sex profoundly shapes brain function, yet its precise influence on neural oscillations was poorly understood. Despite decades of research, studies investigating sex-based variations in electroencephalographic (EEG) signals have yielded inconsistent findings that obstructs what may be a potentially crucial source of inter-individual variability in brain function. To address this, we analyzed five publicly available resting-state datasets, comprising EEG data (n=445) and iEEG data (n=103). Our results revealed striking age-dependent sex differences: older adults (30-80 years) exhibited robust sex differences, with males showing heightened low alpha (8-9 Hz) activity in temporal regions and attenuated low beta (16-20 Hz) oscillations in parietal-occipital areas compared to females. Intriguingly, these sex-specific patterns were absent in younger adults (20-30 years), suggesting a complex interplay between sex and aging in shaping brain dynamics. Furthermore, we identified consistent sex-related activity in the precentral gyrus with the results of scalp EEG, potentially driving the observed scalp EEG differences. This multi-level analysis allowed us to bridge the gap between cortical and scalp-level observations, providing a more comprehensive picture of sex-related neural dynamics. To further investigate the functional implications of these oscillatory differences, we conducted correlation analyses to uncover significant associations between sex-specific oscillatory patterns and several lifestyle factors (behavioral and anthropometric measures) in older adults. This comprehensive investigation demonstrates the complex interplay between sex, age, and neural oscillations, revealing the variability in brain dynamics. And our findings highlight the importance of careful demographic consideration in EEG research design to ensure fairness in capturing the full spectrum of neurophysiological diversity.

**Significance statement:** The influence of biological sex and age on neural oscillations had been a long-standing, unresolved question in EEG research, largely unaddressed due to limited sample sizes and simplistic demographic matching. Our study leverages large-scale, open datasets to tackle this issue, analyzing hundreds of participants across five datasets. Our findings demonstrate substantial sex-based differences in even resting-state EEG baselines, particularly in low alpha and low beta bands, uncovering a significant source of variability in neural activity. By connecting these sex and age-related variations to potential neural circuit mechanisms and lifestyle factors, our findings highlight the importance of careful demographic consideration in EEG research design in EEG experimental design to accurately capture the rich spectrum of neurophysiological variability across the lifespan.

## Introduction

Electroencephalography (EEG) offers unique advantages in recording spontaneous and task-related neural activities due to its high temporal resolution and non-invasive nature (1, 2). It has been used widely in measuring cognitive functions such as attention(3–5), memory(6–8), and learning(9, 10), as well as to investigate neurological and psychiatric conditions including attention deficit hyperactivity disorder (ADHD)(11–13), autism(14–16), depression(17–19), Alzheimer’s disease (AD)(20–22), schizophrenia(23, 24). However, the impact of demographic factors, particularly sex and age, on neural oscillations remains poorly characterized. This gap potentially undermines the interpretation of EEG studies comparing different groups, despite efforts to match demographic variables.

EEG records neural oscillations (2, 25), which could reflect the interactions between different types of neuronal populations (26, 27). Sex differences in brain structure and function are well-documented across species (28), with animal studies revealing specific neural circuit mechanisms (29–31). However, their manifestation in EEG oscillatory features and underlying neural mechanisms remain unclear, despite significant sex differences in human brain structural connectivity(32). Previous studies investigating sex differences in EEG oscillations have yielded inconsistent results (33), across delta(34–38), theta(34, 35, 37, 38), alpha(34, 35, 37–40) and beta(34, 36–39, 41) frequency bands, likely due to limited sample sizes, age confounds, and unclear behavioral correlates. Age itself also significantly affects neural responses(35, 42, 43), with a consistent increase of beta-band activities (44–49). However, the interaction between age and sex differences in EEG oscillations remains poorly understood, as studies examining combined age and sex effects have been limited by small sample sizes (41, 50, 51). This limitation has precluded a comprehensive understanding of how sex differences in neural oscillations might evolve across the lifespan and how they relate to cognitive functions and behavior.

To address these critical gaps, we analyzed large-scale publicly available resting-state datasets (4 EEG, 1 iEEG) to systematically investigate sex differences in neural oscillations across the lifespan. Our approach comprised four key steps: (1) identifying age-dependent sex differences in specific brain regions and frequency bands using a primary EEG dataset (n=203); (2) validating these findings across three additional EEG datasets (n=242 total); (3) corroborating surface EEG findings with deep brain recordings using an iEEG dataset (n=103); and (4) exploring relationships between sex and age -specific oscillatory features and behavioral and anthropometric measures. This comprehensive analysis provides a robust characterization of sex differences in neural oscillations, their evolution with age, and their potential functional significance. Our findings have implications for improving EEG-based diagnostics, and enhancing the design and interpretation of cognitive neuroscience studies.

## Materials and Methods

### Public EEG dataset - Max Planck Institute (MPI) Leipzig Mind-Brain-Body Dataset**(52)**

The EEG data was collected from 203 participants (age from 20 to 80, 129 Male, 74 Female) during resting state (eye-open (EO) and eye-closed (EC)). The public dataset could be download in the following link: https://fcon_1000.projects.nitrc.org/indi/retro/MPI_LEMON.html. The public dataset has been preprocessed; the details could also be seen in the above link. The EEG was recorded with 61 electrodes and the total time length of open- and closed- eye state is 8 minutes. We divided the MPI dataset into two groups: young age (YA, aged 20-30, n=125. 89 Male, 36 Female) and older age (OA, age 30-80, n=78, 40 Male, 38 Female) group.

### Public iEEG dataset - The Montreal Neurological Institute (MNI) iEEG Dataset**(53)**

The iEEG data was collected from 1772 channels with normal brain activity (n=106, age from 13 to 62, 58 Male, 48 Female) during resting state (eye-closed (EC)). The public dataset could be download in the following link: https://mni-open-ieegatlas.research.mcgill.ca. Total time length of closed- eye state is 1 minute. We divided the MNI dataset into two groups: young age (aged <30) and older age (age >=30) group. There are 38 brain regions in total, but size of data in some regions is not sufficient to compare the statistical difference. The criteria to select the brain region for sex difference analysis is that in each age group, for a specific brain region, it should have at least 10 electrodes’ data. Under this criterion, 15 brain regions were selected in OA group, 9 brain regions were selected in YA group.

### Public EEG dataset - Southwest University (SU) Dataset**(54)**

The EEG data was collected from 60 participants (age from 18 to 28, 28 Male, 32 Female) during resting state (eye-open (EO) and eye-closed (EC)). The public dataset could be download in the following link: https://openneuro.org/ datasets/ds004148/versions/1.0.1. The public dataset has been preprocessed; the details could also be seen in the link. Three participants were excluded due to their data is incomplete (3 Female). The EEG was recorded with 61 electrodes and the total time length of open- and closed- eye state is 9 minutes.

### Public EEG dataset - Healthy Brain Network (HBN) Dataset**(55)**

The EEG data was collected from 71 participants (age from 18 to 22, 39 Male, 32 Female) during resting state (eye-open (EO) and eye-closed (EC)), for the subjects who is younger than 18 is not considered in this study. The public dataset could be download in the following link: https://openneuro.org/ datasets/ds004186/versions/2.0.0. The public dataset has been preprocessed; the details could also be seen in the link. The EEG was recorded with 128 electrodes and the total time length of closed- eye state is 40 seconds.

### Public EEG dataset - SRM Dataset**(56)**

The EEG dataset comprises resting-state recordings from 111 participants (age from 17 to 71, 42 Male, 69 Female) during eye-closed (EC) conditions. Data were collected using a 64-electrode montage for a total duration of 4 minutes per participant. The preprocessed dataset is publicly available at https://openneuro.org/datasets/ds003775/versions/1.2.1/download. We divided the MNI dataset into two groups: young age (aged <30) and older age (age >=30) group. Detailed preprocessing steps were provided in the dataset documentation.

### Data Analysis

Data processing was performed in MATLAB (www.mathworks.com) with custom scripts. Data in three dataset was filtered between 1 and 30 Hz. The reference of MPI dataset and SU dataset was set to the averaged reference. We used spectrum analysis to quantify the neural oscillation strength in all electrodes in open or closed-eye state. The power spectral density (PSD) for time series EEG and iEEG data each electrode was computed using the multi- taper method with 5 tapers using the Chronux toolbox(57), which is an open-source, data analysis toolbox available at http://chronux.org. Similar methods have been applied in various biomedical fields, such as experimental neuroscience(58–61), neuropsychological disorders(12, 14, 24), etc. Relative power was calculated by dividing the power at a specific frequency by the total power summation from 3 to 30Hz.

### Statistical Analysis

The pairwise t-test was conducted to examine the relative power in alpha-band (8-12 Hz) between EO and EC state in MPI dataset (Fig 1C) and SU dataset (Fig 3B). In MPI dataset, in order to screen out significant frequency bands and electrode positions for sex difference, the independent t-test was tested difference between relative power of male and females in YA and OA group in open and closed-eye state respectively (Fig 1D, similar statistics also conducted in Fig 3C in SU Dataset). Then, two-way ANOVA (gender (M and F) and age (OA and YA)) was conducted to relative power in low alpha (LA), low beta (LB) band in open and closed- eye state respectively. Further, we used t-test for pairwise comparison (Fig 2EF). In MNI Dataset, t-test was conducted to test the significance of sex difference of relative power in LA and LB band each brain region for YA group and OA group respectively.

**Figure 1.**
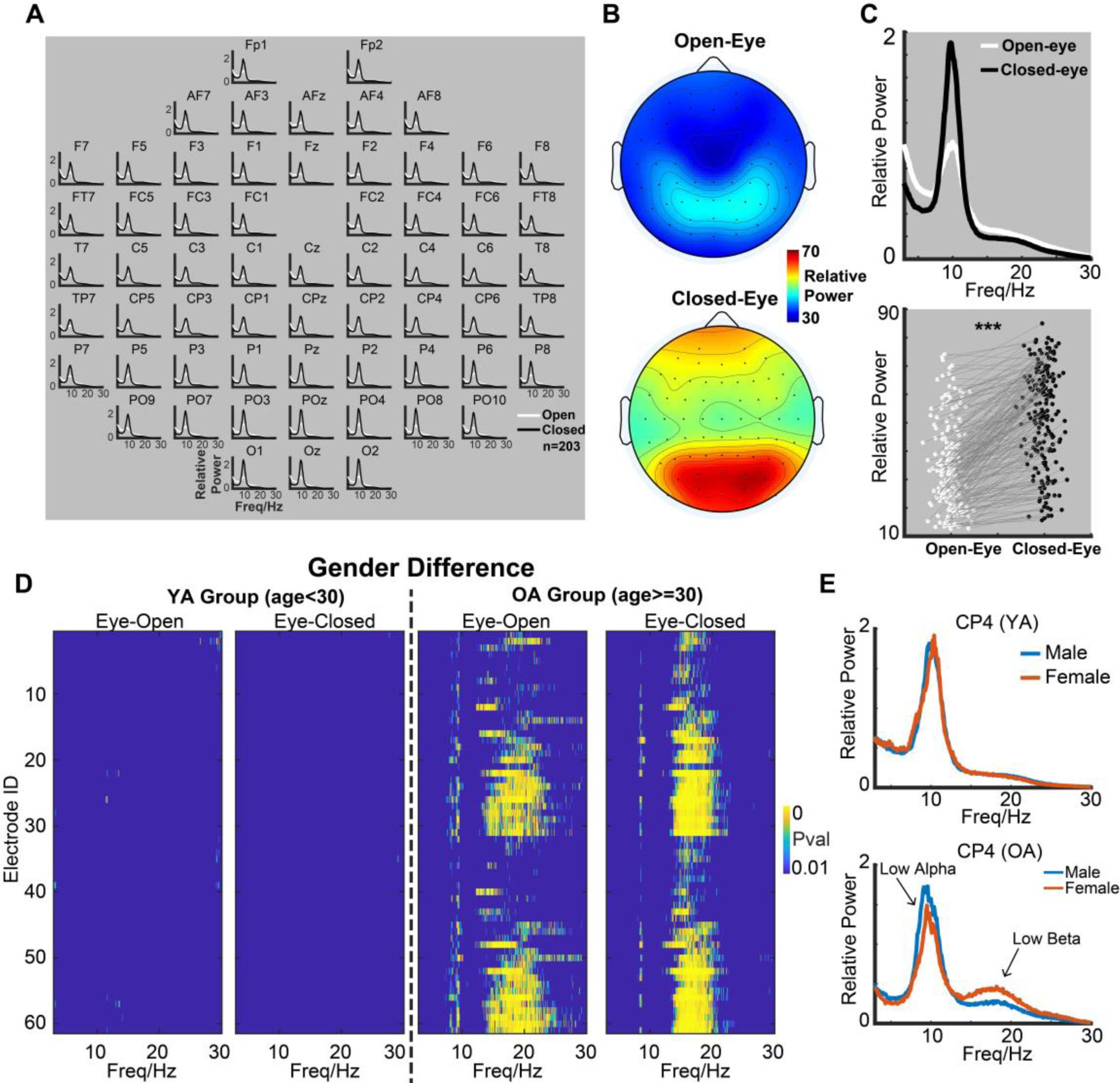
Age-dependent sex difference in low alpha and low beta EEG oscillations. A. Averaged power spectrum (n=203) across 61 electrodes in MPI dataset during eyes-open (white curve) and closed (black curve) states. B. Average topographic map of alpha (8-12Hz) power in eyes-open and eyes- closed states. C. Upper: Grand average power spectrum across all electrodes. Lower: Comparison of alpha power (8-12 Hz) between eyes-open and eyes-closed states. Each dot represents an individual subject. D. Statistical significance of sex differences in the eyes-open and eyes-closed states for young adult (YA, left) and older adult (OA, right) groups. X-axis is frequency and y-axis is electrode ID. E. Power spectrum of male (blue curve) and female (red curve) in closed (black curve) state in a typical example electrode (CP4) for demo in YA (above panel) and OA (below panel) groups.

**Figure 2.**
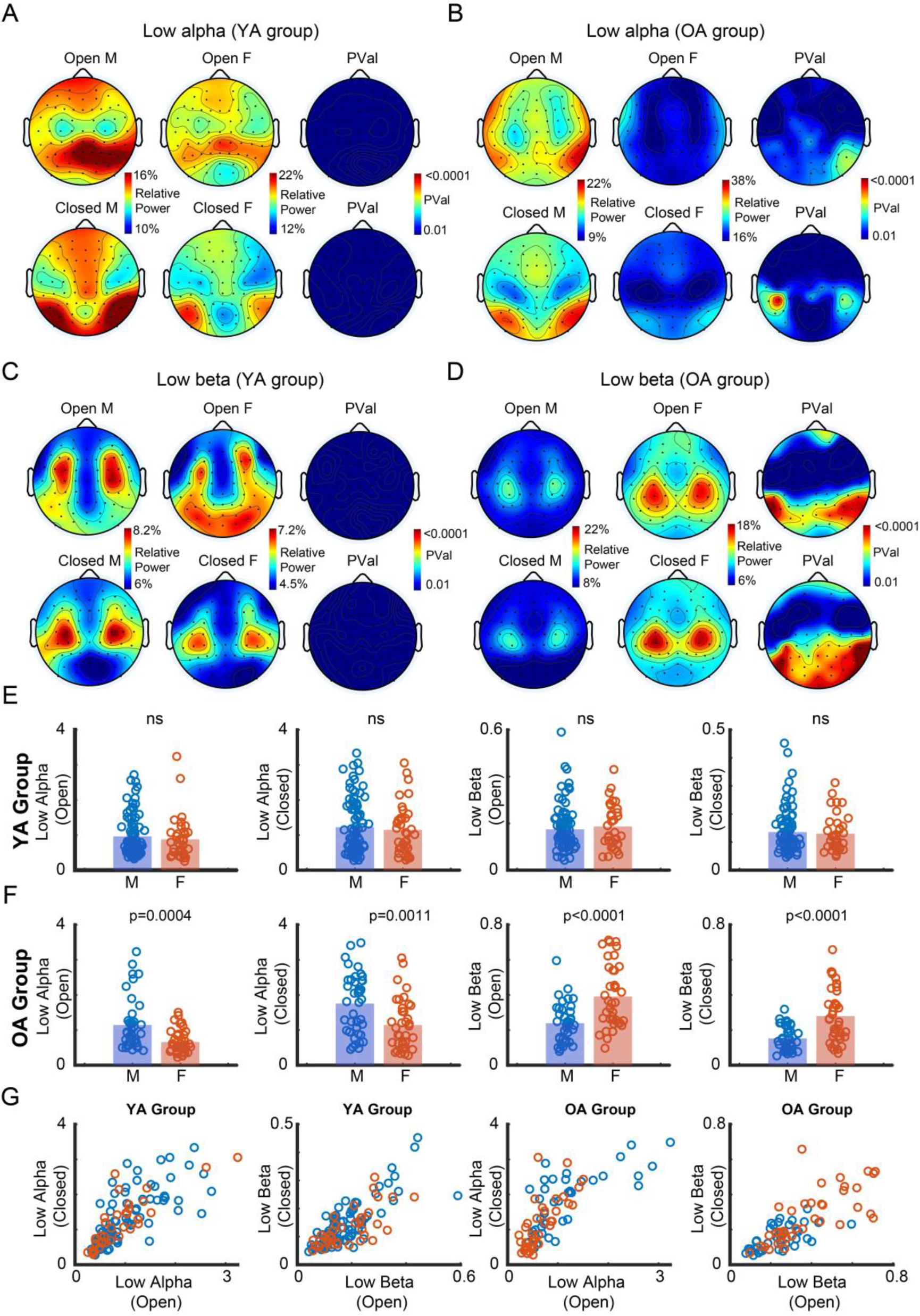
Sex difference with distinct topographic maps in low alpha and low beta band. (A-D) Topographic maps of EEG power for males (left), females (middle), and sex difference significance (right; red: significant, blue: non-significant) in eyes-open (top) and eyes-closed (bottom) states. (A) Low alpha, young adults (YA). (B) Low alpha, older adults (OA). (C) Low beta, YA. (D) Low beta, OA. E. Sex differences in low alpha and low beta power for YA in eyes-open and eyes-closed states. F. Sex differences in low alpha and low beta power for OA in eyes-open and eyes-closed states. G. Scatter plots comparing eyes-open versus eyes-closed power in low alpha and low beta bands for YA and OA. Each dot in (E-G) represents an individual subject.

**Figure 3.**
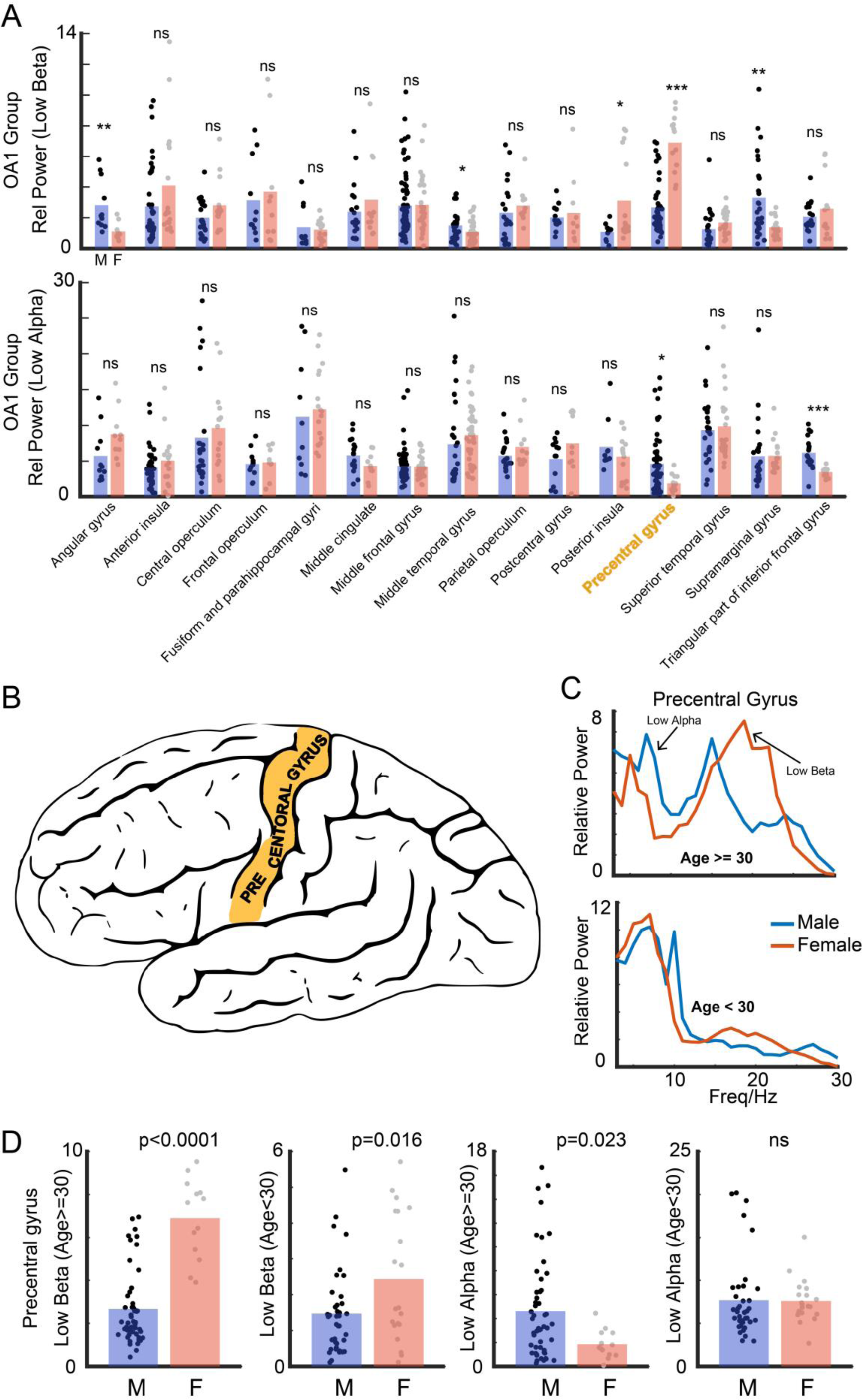
Sex differences in precentral gyrus neural responses align with EEG findings. A. Comparison of 15 brain regions in iEEG dataset (age >=30) in older adults group in closed-eye state in low beta (above) and low alpha band (below), where * is for p<0.05, ** is for p<0.01, *** is for p<0.001, and ns is for not significant. B. Illustration of anatomical location of precentral gyrus. C. LFP power spectra in precentral gyrus for younger (age<30) and older (age>=30) adults, by sex (male: blue; female: orange). D. Sex differences in low alpha and low beta power in precentral gyrus for younger and older adults. Each dot represents a recording site.

## Results

Analysis of the MPI dataset revealed a robust increase in alpha-band activity during eyes-closed (EC) compared to eyes-open (EO) conditions across most recording sites (Fig. 1A). This effect was most pronounced in the parietal-occipital region (Fig. 1B). By averaging all electrodes, most subjects showed significant (p<0.0001) increase in the alpha power (Fig 1C).

### Age dependent sex difference on low alpha- and low beta- band activities

The dataset was separated with two age groups (OA and YA, see methods), and we compared the sex difference from 3 to 30Hz in all electrodes in EO and EC state (Fig 1D). In the YA group, no significant sex differences in relative power were observed across all electrodes in both eyes-open (EO) and eyes-closed (EC) states (Fig. 1D, left panels). In contrast, the OA group exhibited significant sex differences (p<0.01) in multiple recording sites, specifically in the low alpha (LA, 8-9 Hz) and low beta (LB, 16-20 Hz) bands, during both EC and EO states (Fig. 1D, right panels). Taking CP4 site as an example (Fig 1E), while spectral profiles of males and females in the YA group were indistinguishable, the OA group displayed a clear divergence: males exhibited higher LA power, whereas females showed greater LB power.

### Sex difference with distinct topographic maps in low alpha and low beta band

Based on the statistical analysis as sown in Fig. 1D, subsequent investigations focused on the low alpha (LA, 8-9 Hz) and low beta (LB, 16-20 Hz) bands. Topographic mapping of sex differences revealed age-dependent patterns (Fig. 2A-D). In the young adult (YA) group, no significant sex differences were observed in either LA (Fig. 2A) or LB (Fig. 2C) bands. Conversely, the older adult (OA) group exhibited significant sex differences (p<0.01) in parietal-temporal regions for LA (Fig. 2B) and parietal-occipital regions for LB (Fig. 2D). Two-way ANOVA (age and sex as factors) on the five most significant electrodes for each condition (EO or EC in LA or LB) revealed significant main effects and interactions (p<0.001). The pairwised comparisons are shown in Fig 2E for YA group, and Fig 2F for OA group. From the scatter plot (Fig 2G), based on the 2-dim oscillatory features (EC and EO) in LA and LB, we could find clear separation between males and females in OA group.

To assess the generalizability of these findings, analyses were extended to three additional open EEG datasets (SU, HBN, and SRM; Fig. S1). The predominantly young adult SU and HBN datasets showed no significant sex differences in LA or LB bands (Fig. S1, top two rows). However, the age-diverse SRM dataset corroborated the MPI findings, exhibiting similar sex-related trends in both LA and LB bands (Fig. S1, bottom row).

### Consistence for sex difference of neural response in precentral gyrus with EEG finding

To further explore the potential neural circuits contributes to the findings in MPI dataset, analyses were extended to an intracranial EEG (iEEG) dataset (MNI dataset; Fig. 3A).. In the older adult (OA) group, significant sex differences (p<0.05) were observed in multiple brain regions. Females exhibited higher low beta (LB) power in the posterior insula and precentral gyrus, while males showed higher low alpha (LA) power in the precentral gyrus and inferior frontal gyrus. Additionally, males displayed significantly higher LB power (p<0.01) in the angular gyrus and supramarginal gyrus.

Notably, the spectral pattern of sex differences in the precentral gyrus (Fig. 3B, C) closely mirrored the scalp EEG findings from the MPI dataset. Age-stratified analyses (Fig. 3D) revealed that in the LB band, OA females exhibited significantly stronger relative power compared to males (p<0.0001), with this effect attenuated in the young adult (YA) group (p=0.016). Conversely, in the LA band, OA males showed significantly stronger relative power than females (p=0.023), while no significant difference was observed in the YA group.

### Potential lifestyle-related factors associated with the sex difference in low alpha and low beta activities during aging

To investigate the potential factors that may relate to the sex difference in LA and LB band in MPI dataset, anthropometries (height, weight, waist and hip) were considered to test for the relationship (Fig 4 A-B). In the older adult (OA) group, only females exhibited significant positive correlations between LB power and hip circumference (p<0.01; Fig. 4A-B). Similarly, body mass index (BMI), derived from anthropometric data, showed a significant positive correlation with LB power exclusively in OA females (p<0.01; Fig. 4C-D).

**Figure 4.**
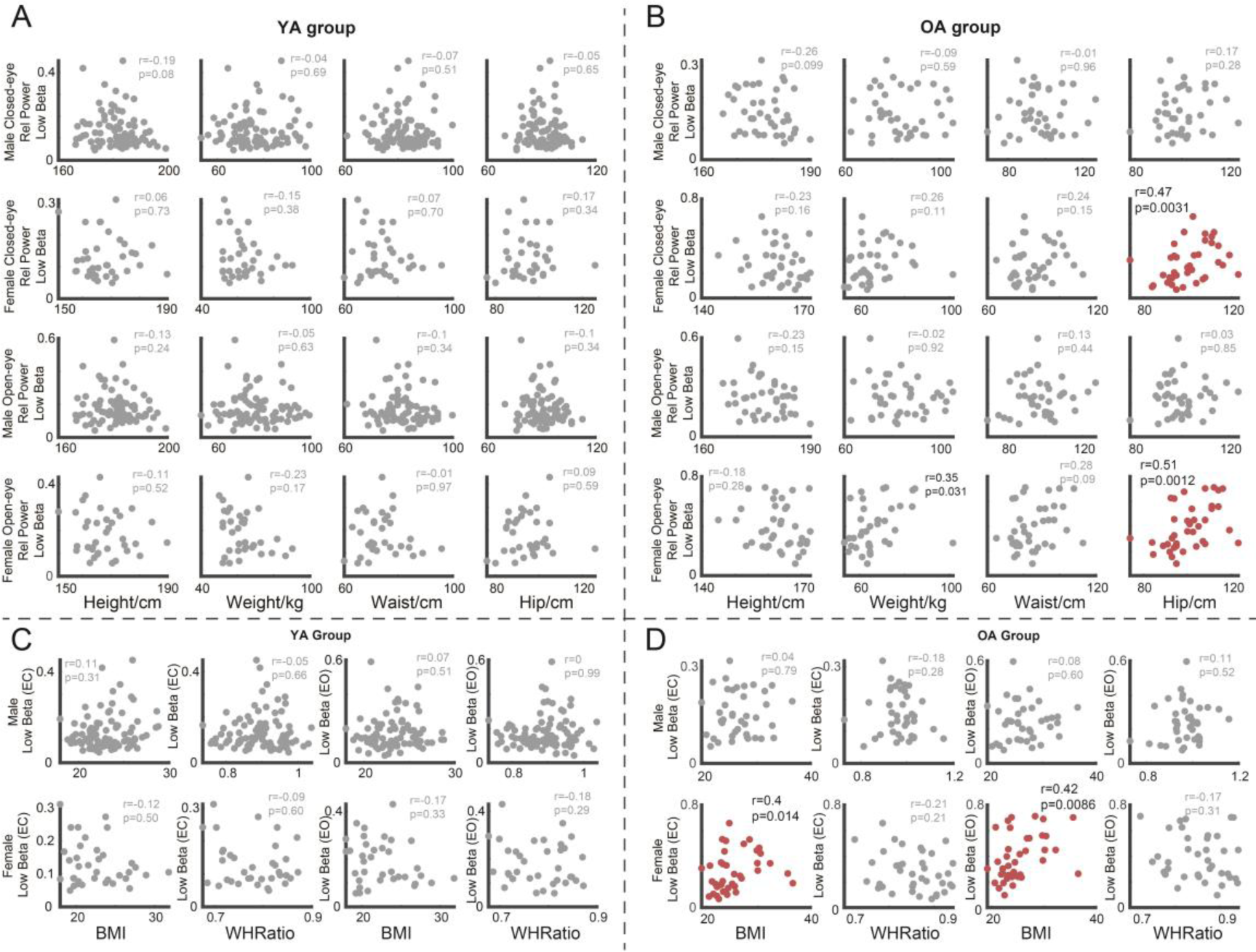
Associations between anthropometries and low beta activity. (A, B) Relationships between low beta power and height, weight, waist, and hip measurements in (A) young adults (YA) and (B) older adults (OA) during eyes-open and eyes-closed states. (C, D) Relationships between low beta power and BMI and waist-hip ratio in (C) YA and (D) OA during eyes-open and eyes-closed states.

Further analyses investigated potential links with mood and alcohol consumption using the Hamilton scale and alcohol-related scores (Fig. S2 A-B). While no significant correlations were found between Hamilton scale scores and LA or LB power in either age group, alcohol-related scores showed significant associations with both LA (positive correlation) and LB (negative correlation) power across eyes-closed and eyes-open states (all p<0.05).

## Discussion

In this study, we have, for the first time, provided novel insights into sex differences in neural oscillations across lifespan and identified some potential factors contributing to these differences by analyzing five public datasets encompassing both scalp and intracranial EEG recordings. Our findings reveal age-dependent sexual dimorphism in neural oscillatory patterns, with older males exhibiting enhanced low alpha (LA) power in temporal-parietal regions, while older females show increased low beta (LB) power in parietal-occipital areas. In particular, while older females show increased low beta (LB) power in parietal-occipital areas. Notably, these scalp-level observations are corroborated by intracranial recordings from the precentral gyrus, suggesting a potential neural substrate for the observed sex differences. Furthermore, we also found that in older adults, sex-specific differences in LA and LB band power are significantly associated with alcohol consumption patterns, while the LB band activity is significantly correlated with hip circumference in aging females. These findings collectively highlight the complex interplay between aging, sex, and lifestyle factors in shaping neural oscillatory patterns, potentially offering new avenues for understanding sex-specific trajectories of cognitive aging and their modulation by environmental influences.

### Mechanisms underlying age-dependent sex difference in neural oscillations

Our study advances the field beyond previous investigations of sex differences in EEG (33, 39) by integrating age as a critical variable, validating findings with intracranial EEG data, and exploring correlations with behavioral and psychometric measures. Our study is of significant scientific importance, particularly in highlighting that sex and age should be key variables to control for in future EEG experiments. In our study, sex differences in EC or EO states are similar (Fig 1D, Fig 3C), indicating that the generality of sex differences may not be task-specific.

The contribution of sex differences in aging may not be attributed to a single brain region, but likely involves the coordinated action of multiple brain regions. For example, while the precentral gyrus showed high concordance with scalp EEG findings in both low alpha (LA) and low beta (LB) bands, other regions also exhibited significant sex differences (Fig. 3). The distinct spatial patterns of significance for LA and LB (Fig. 2B, 2D) align with proposed divergent mechanisms underlying alpha(62, 63) and beta(64) oscillations. Enhanced LB power in females and LA power in males each implicate multiple brain regions, highlighting the distributed nature of these sex differences. Furthermore, our analyses identified two brain regions where males exhibit higher low beta band power compared to females, a pattern not apparent in the broader topographical assessment.

### Associations between daily behaviors and neural oscillations

Neuronal oscillations underpin diverse cognitive functions(5, 61, 65), and are intricately connected to human behaviors (15, 66, 67). Our analyses reveal significant associations between lifestyle factors and oscillatory power in older adults. Notably, we observed an inverse relationship between alcohol-related scores and the power of low alpha (LA) and low beta (LB) bands. Higher alcohol-related scores correlated with increased LA and decreased LB power. While causality cannot be inferred, this association, considered alongside alcohol’s known health effects and GABAergic modulation(68), suggests that neural oscillatory patterns might serve as potential indicators of recent alcohol consumption.

Our investigation into anthropometric correlates uncovered a positive association between LB power and both hip circumference and BMI in older females. Meanwhile, waist-hip ratio (69, 70) showed no significant correlation (Fig 6 C-D). The relationship between LB power and hip circumference may reflect complex interactions between body composition and neural activity, potentially influenced by age-related changes in motor function (49).

These correlational findings highlight the intricate relationships between lifestyle factors, body composition, and neural oscillations in aging populations. While causal relationships remain to be established, these associations suggest that EEG-derived measures might provide insights into lifestyle-related variations in brain function. This non-invasive approach warrants further investigation for its potential in health monitoring and intervention strategies, particularly in aging populations where lifestyle factors are known to impact cognitive health.

### Limitations and future work

While our findings demonstrate consistent sex differences in neural oscillations across diverse datasets, the use of publicly available data from various geographical origins introduces potential confounds related to racial and ethnic factors. Although the consistency of our results suggests these factors may not substantially influence the observed sex differences, they likely contribute to some variability. Future investigations should systematically control for racial and ethnic factors to refine our understanding of sex-specific neural dynamics across populations. Longitudinal studies tracking individuals across the lifespan, combined with multimodal neuroimaging techniques, would elucidate the developmental trajectories of these sex differences and their underlying neuroanatomical substrates.

## Acknowledgments

This work was funded by the Research Grants Council of the Hong Kong Special Administrative Region, China (Project No. R4022-18, N_CUHK456/21, 14114721, and 14119022 to V.C.K.C. Project No. 9042986 to R.H.M.C.), and funding from City University of Hong Kong (No. 7005641 to R.H.M.C.)

## Conflicts of interest statement

The co-authors declare that the research was conducted in the absence of any commercial or financial relationships that could be construed as a potential conflict of interest.

## Contributors

CH, RC and VC conceived and designed the study. CH contributed to the literature search, data analysis, and the interpretation of results. All authors contributed to writing the paper.

---

**Supplementary Figure 1.**
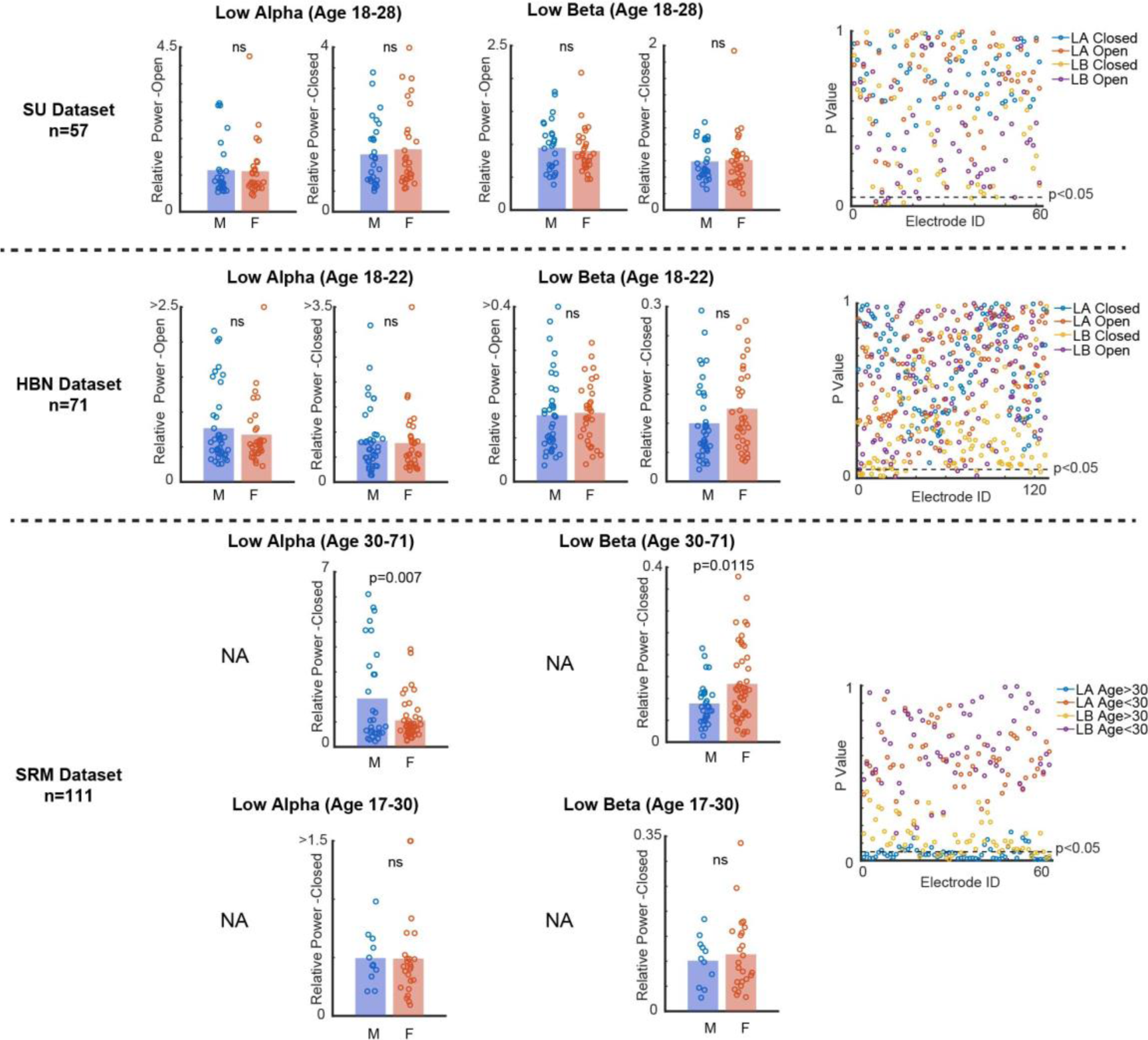
Consistency of sex differences in low alpha and low beta EEG activity across additional datasets. Left panels: Sex differences in low alpha (LA) and low beta (LB) power during eyes-open and eyes-closed states for SU, HBN (primarily younger adults), and SRM (both younger and older adults) datasets. Each dot represents an individual subject; bars indicate sample means. Right panels: Electrode-wise p values for sex differences. In younger adults, most electrodes show insignificant differences. In older adults, LA and LB exhibit significant differences across multiple electrodes.

**Supplementary figure 2.**
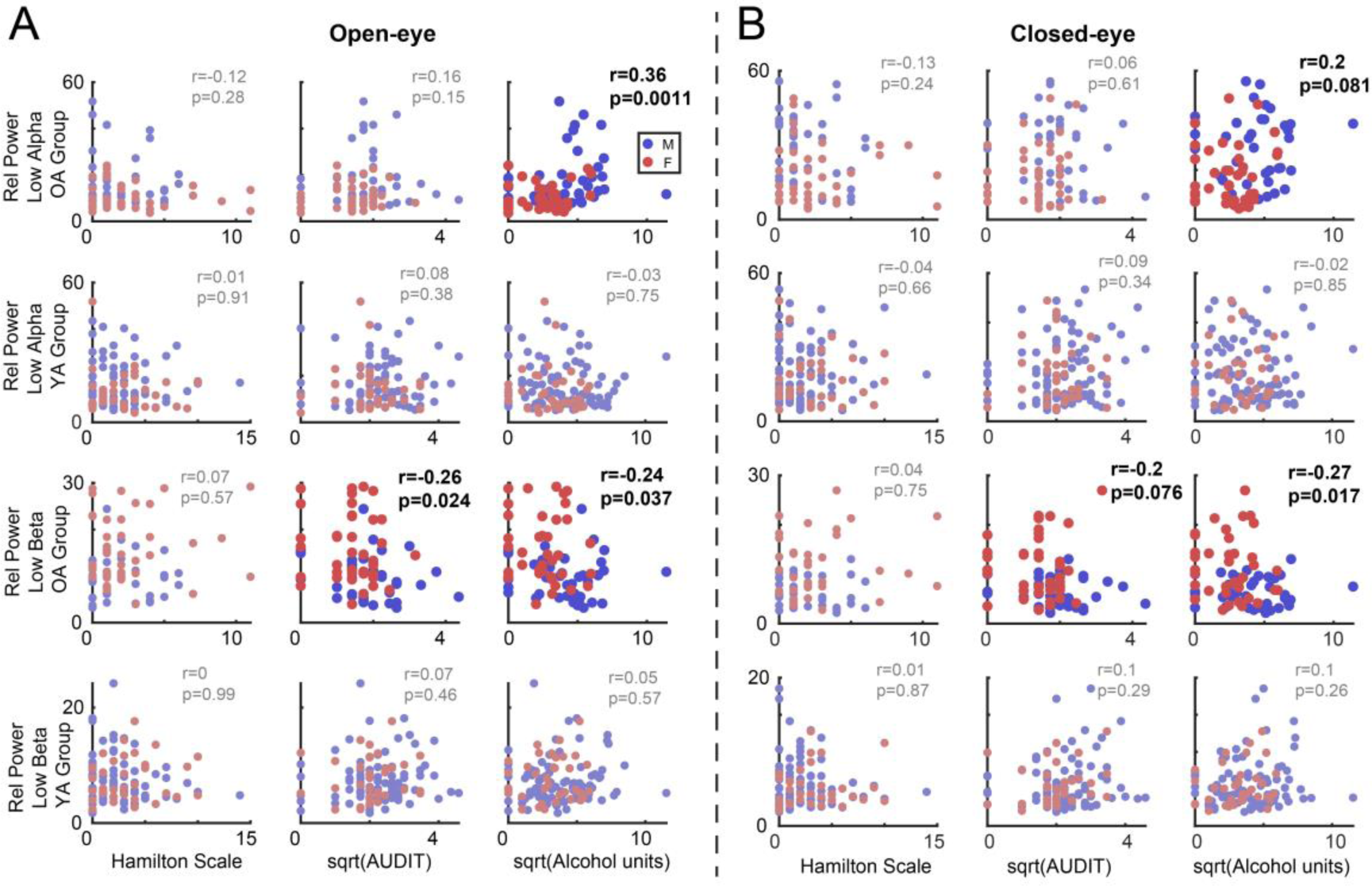
Associations between lifestyle factors and neural oscillations. A. Scatter plot of scores (Hamilton, AUDIT, Alcohol units) and relative power (low alpha and low beta) during eyes-open state. B. Scatter plot of scores (Hamilton, AUDIT, Alcohol units) and relative power (low alpha and low beta) during eyes-closed state. Each dot represents an individual subject.

## Notes

### Competing Interest Statement

The authors have declared no competing interest.

